# Fab-arm exchange affects each and all endogenous serum IgG4 evenly

**DOI:** 10.1101/2025.03.27.645639

**Authors:** Linus Wollenweber, Albert Bondt, Maartje G. Huijbers, Albert J.R. Heck

## Abstract

Antibodies are key molecular elements of the human immune system and account for an increasingly large proportion of therapeutics. Next to the well-studied and explored IgG1, several other classes (e.g. IgA, IgM) and sub-classes (e.g., IgG2, IgG3 and IgG4) exist in humans. In particular IgG4 is worth a closer examination as it has unique natural properties and is regularly used as a scaffold in biologicals. IgG4 stands out from the other IgG by its ability to dissociate and form two half-molecules which can interchange with those of other clones to form novel bivalent antibodies. Detailed analysis of endogenous IgG4 so far has been hampered by a lack of analytical methods to dissect and analyze IgG4 molecules with clonal resolution. Here, we present an LC-MS-based approach enabling the analysis of IgG4 repertoires with clonal resolution, which we used to monitor endogenous serum IgG4 repertoires from seven healthy donors. Most strikingly, our data reveal the combinatorial explosion in diversity of the serum IgG4 clonal repertoire. This phenomenon is explained by the stochastic behavior of Fab-arm exchange, making virtually each IgG4 molecule in serum bispecific. Although the endogenous IgG4 clonal repertoire is therefore extremely diverse, we demonstrate that this IgG4 repertoire persists over time within an individual healthy donor for more than one year. This newly established method now enables repertoire analysis for IgG4, which plays a critical role in a plethora of disease settings including allergy, autoimmunity and vaccination.

## Introduction

It is tempting to consider the four subclasses of IgG (IgG1, IgG2, IgG3, IgG4) as similar, as they share many structural properties and show over 90 % amino acid sequence homology. However, as often in biology, the devil is in the details and there are many subtle but important differences between subclasses of IgG, affecting both their structure and function^1,2^. These include among others differences in light to heavy chain coupling *via* disulfide bonds (IgG1 vs IgG2-4)^2^, length and flexibility of the hinge region^3^, and post-translational modifications they can harbor^4,5^.

The IgG4 subclass is commonly described as the odd one out among the other IgG^6,7^. At first glance, this cannot be explained by its generic structure (Figure 1A), as all four IgG subclasses are similar in that regard, being composed of two pairs of heavy (Hc) and light chain (Lc). IgG4 stands out from the other IgG sub-classes due to its relatively labile Hc-Hc interactions, allowing IgG4 antibodies to dissociate rather easily in two half-molecules (Figure 1B). Consequently, an exchange can occur *in vivo* and *in vitro*, in which half-molecules of different IgG4 antibody molecules interchange to form novel bivalent antibodies (Figure 1B). This so-called Fab-arm exchange (FAE) process, being in humans unique for IgG4, has been known for some time^8-16^, but key questions regarding the relative proportion of non-covalently linked IgG4 half-molecules or the extend of the Fab-arm exchange *in vivo*, e.g. in blood, have not yet been fully addressed. A major bottle-neck in addressing these questions is a lack of methodology to dissect and analyze simultaneously all IgG4 molecules present in a body fluid with clonal resolution, basically recording the secreted IgG4 antibody repertoire.

**Figure 1.**
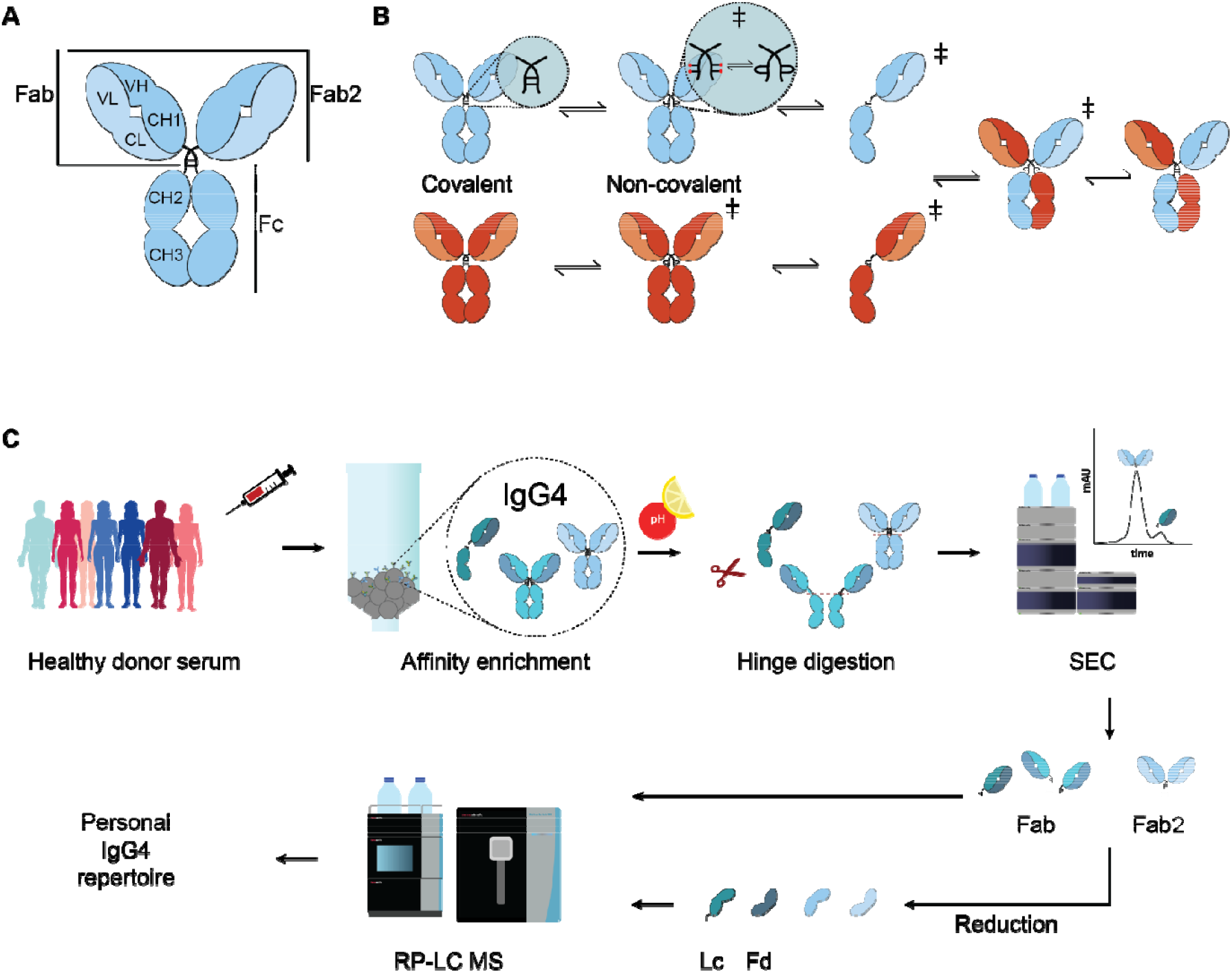
Structural features of IgG4 and approach to analyze IgG4 clonal repertoires. A) Generic structure of IgG with used domain and subunit nomenclature. B) IgG4 co-exists in two structural isoforms, namely covalently and non-covalently linked half-molecules. Non-covalently linked half-molecules are comprising molecules with reduced and intrachain hinge disulfides. Under the right circumstances half-molecules in this transition state can dissociate and reassemble with other half-molecules of different sequence and antigen specificity (adopted from Rispens et al.^8^). C) Analyzing IgG4 repertoires is initiated by the selective enrichment of serum IgG4 of seven individual donors. Purified IgG4 is subsequently digested at a specific site in the hinge region, releasing IgG4 Fab2 and Fab molecular fragments that contain still all information on the hypervariable CDRs and the Fd-Lc coupling. Next, SEC is used to separate the population of IgG4 Fab2 (∼98 kDa) and Fab (∼49 kDa) molecules. Fab and Fab2 clones are separated and both fractions are mass analyzed by LC-MS either intact or upon further reduction of these fragments to generate and analyze their Lc and Fd fragments.

A better knowledge on the nature and extend of Fab-arm exchange *in vivo* is not purely of fundamental immunological interest but may have also important clinical implications, as IgG4 represents the second most used platform for antibody therapeutics. An IgG4 format is often selected in therapeutics when effector functions are undesirable, for example when a mAb is mostly used to block a specific target as in immune checkpoint therapies^2,17-19^. The low level of effector functions is one of the unique characteristics of IgG4, likely in place to dampen an overshooting immune response following an allergy^20,21^ or parasitic infection^22^.

Here, we present a liquid chromatography mass spectrometry (LC-MS)-based approach geared at the selective clonal profiling of endogenous IgG4 as present in body fluids (Figure 1C). Our approach starts with an enrichment of the IgG4 population from a limited amount of sample (∼100 µL of serum). Subsequently, all IgG4 molecules are cleaved using the hinge-directed protease IdeS, releasing the IgG4 (Fab2) domains, thereby importantly retaining the cognate VH-VL pairing and complementarity determining regions (CDR). Following digestion, we observed that the resulting eluate contained both Fab2 and Fab fragments. Size-exclusion chromatography (SEC) allowed us to directly compare the IgG4 repertoire compositions of the initially pool of covalently and non-covalently linked IgG4s. Cumulatively, our data provides solid evidence that Fab-arm exchange is a process occurring stochastically *in vivo*, affecting all endogenous IgG4s evenly. As a result, IgG4 repertoire diversity in serum is massively more extensive compared to that of other IgG subclasses (e.g., roughly from n for IgG1 to 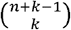 for IgG4).

## Results

### IgG4 hinge digestion uniquely provides two distinct mass populations

Recently, we introduced two distinct mass spectrometry-based approaches to specifically profile IgG1 or IgA1 repertoires (at the protein level) in body fluids with clonal resolution^23,24^. To a certain extent, these approaches are modular, and thus are suitable to be expanded to other classes and sub-classes of immunoglobulins. The main variable modular components are the affinity resins used to purify the specified isotype (IgG or IgA), and the proteases (IgdE/BdpK^25^ and OpeRATOR, for IgG1 and IgA1, respectively) used to cleave subclass specific the Fab-arms from the Fc part, just *above* the Ig hinge region, releasing the Fab domains. Here, we aimed to extend this approach to IgG4. Since there is no protease yet available which specifically cleaves in the hinge of IgG4, we opted for the use of an IgG4-specific affinity matrix. Subsequently, we used the protease IdeS, which cleaves all human IgG subclasses *below* the hinge region, thereby releasing the Fab2 molecules from the IgG’s.

Based on SEC-UV, we validated Fab2 formation by comparing IdeS induced cleavage patterns of all sub-classes of IgG. Therefore, we used recombinant CD20-targeting mAbs (clone 7D8), which were produced as IgG1, IgG2, IgG3 or IgG4, as previously described^26,27^. In sharp contrast to what was observed for the IgG1, IgG2 and IgG3 mAbs that formed exclusively intact Fab2 molecules, IdeS digestion of the IgG4 mAb resulted in the generation of two populations of fragments, namely Fab2 (∼98 kDa) and Fab (∼49 kDa) molecules (see Figure 2A). We assumed this was caused by co-occurrence of the hinge disulfide-bridge isomers in IgG4 (Figure 1B and 2B). To test whether this observation also applied to endogenous antibodies, we employed SEC to assess the population of Fab2 or Fab molecules present following IdeS digestion of all serum IgG4 enriched by the CaptureSelect IgG4 (Hu) affinity matrix. As a control the same procedure was performed for serum IgG1 enriched by CaptureSelect IgG1 (Hu) (Figure 2C). Also, this analysis clearly revealed the co-occurrence of two abundant populations (Fab2 and Fab) for IdeS-generated fragments of IgG4, but only one population (Fab2) for IgG1 fragments. Moreover, we also observed substantial fractions of both Fab2 and Fab molecules when analyzing two other IgG4 mAbs (Natalizumab and DNP-G2a2-IgG4; Figure2D).

**Figure 2.**
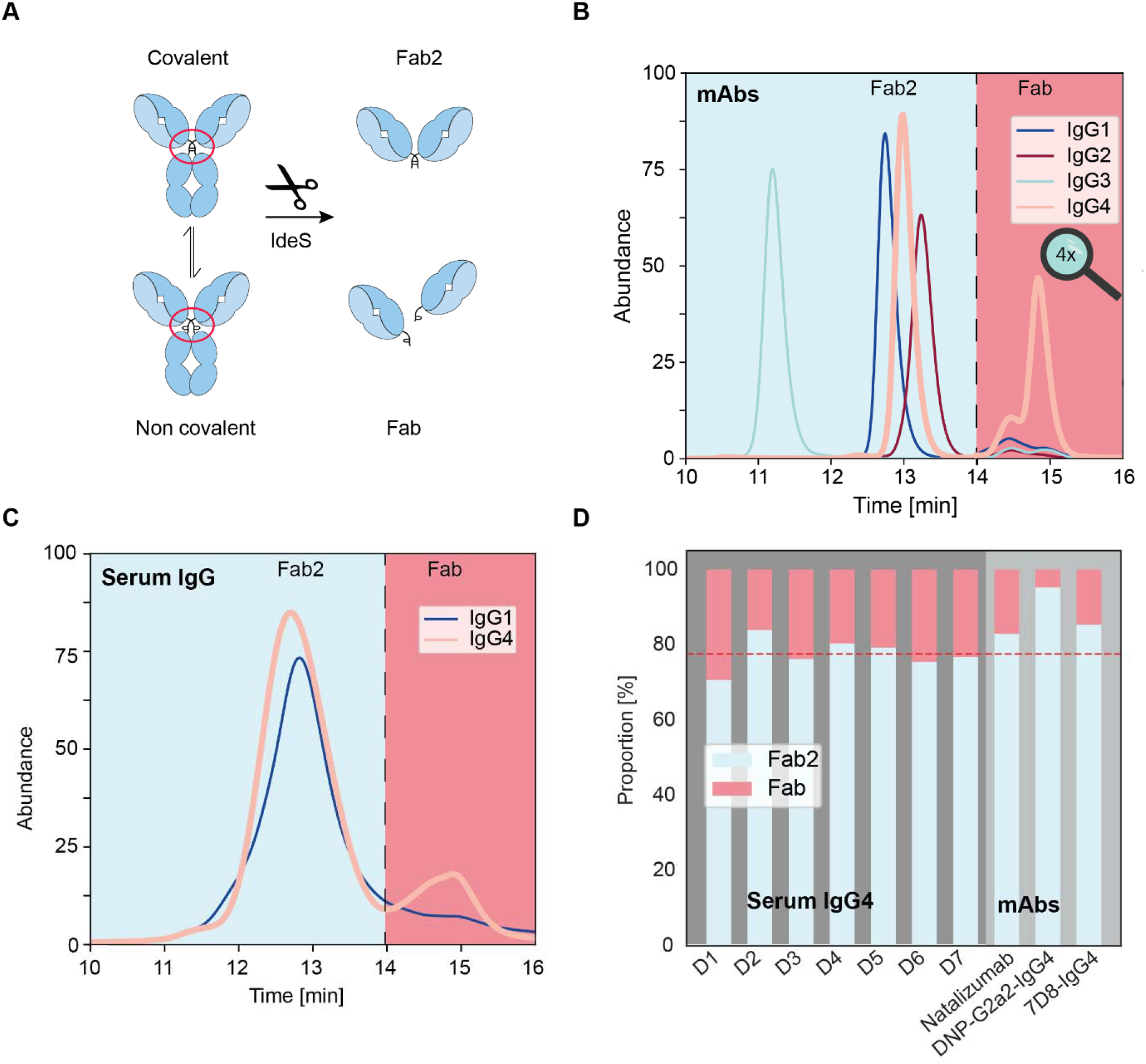
Analysis of IgG4 structural isomers by SEC-UV. A) The unique IgG4 behavior can be traced back to its hinge disulfide structural isomers which lay the foundation for two different products upon IdeS hinge digestion. The covalently bound IgG4 isomer leads to Fab2 molecules and the non-covalently bound IgG4 to Fab molecules. B) SEC-UV traces following hinge digestion of monoclonal IgG1 (dark-blue), IgG2 (dark-red), IgG3 (cyan) and IgG4 (pink). The background color indicates the corresponding products, Fab2 (light-blue) and Fab (red). IgG4 is the only IgG generating two distinct products upon IdeS induced digestion. All these four 7D8 monoclonals are directed against the same epitope and differ only in the subclass specific regions. C) The co-occurrence of (Fab2 and Fab) fragments is also observed for endogenous serum IgG4. The plot layout is the same as in A) without the right y-axis being magnified. D) Stacked bar plots displaying the proportions of Fab2 (light-blue) and Fab (red) as found in seven healthy donors (dark-grey panel) and in three analyzed IgG4 mAbs (light-grey panel). The horizontal red dashed line indicates the average proportion of Fab2 in serum IgG4 (77.6 %) and lies close to the Fab2 proportions found across all donors (SD 3.9 %). In contrast, the investigated mAbs display a higher proportion of Fab2 and a wider spread (SD 5.4 %).

### The proportional co-occurrence of IgG4 structural isomers appear similar between healthy individuals

Having established that serum IgG4 co-occurs in two hinge-disulfide structural variants, we aimed to analyze full IgG4 clonal repertoires in individual donors, by subjecting the Fab2 and Fab fragments to intact protein LC-MS. We therefore analyzed a small cohort of volunteering healthy donors (see materials and methods), consisting of 5 females, and 2 males (30 – 60 years). Furthermore, one of these donors (donor 2) donated serum over a course of 16 months at seven different time points, allowing us to perform longitudinal profiling of this donor’s IgG4 repertoire. For each individual serum sample, we first affinity-enriched all serum IgG4, and subjected this whole pool to digestion by IdeS. We next fractionated these digests using SEC and collected the Fab2 and Fab fractions separately. From the SEC data we could estimate the ratio between covalently and non-covalently linked IgG4 molecules in serum, by integrating the areas under the curves in the SEC-UV traces. Across all donors we found a consistent ratio of 4:1 of Fab2:Fab (mean 77.6 % Fab2, SD 3.9 %, range 70.7 to 83.9 %), indicating that in serum the equilibrium is more on the side of covalently linked IgG4 compared to non-covalently linked IgG4 (Figure 2D, and Supplementary Table 1). The three monoclonals analyzed here displayed on average a slightly higher proportion of Fab2 (mean 87.9 %) than endogenous serum IgG4.

### In serum virtually all IgG4 clones are subjected to Fab-arm exchange

Next, we set out to investigate and analyze the clonal repertoires in the SEC separated Fab2 and Fab fractions at the intact mass level (i.e., ∼98 and ∼49 kDa, respectively for Fab2 and Fab) by LC-MS. The LC-MS readout of the Fab fractions allowed us to observe intact repertoires, detecting several hundreds of clones per donor, looking in diversity and abundance alike to what we observed previously in studies focusing on IgG1 (Figure S1A)^23,28^. In sharp contrast, we could barely detect any intact proteins when analyzing the Fab2 fraction by LC-MS, despite injecting similar total amounts of protein (Figure S1B). We considered that this might be due to the higher Mw of the Fab2 molecules (∼98 kDa), compared to the Fab fragments (∼49 kDa), which can hamper detectability in mass spectrometry. However, similar IgG1 samples allowed for detection of hundreds of clones in the Fab2 profile, indicating that size was not the issue (Figure S1, C). To follow the fate of the IgG4 Fab2, we chose to chemically reduce the Fab2 and Fab fractions obtained from the same IgG4 serum samples, thereby breaking all disulfide bridges. This generates solely Lc (∼23 kDa) and the variable domain bearing fragment of the Hc (Fd; ∼25 kDa). Performing this Lc-Fd profiling of the reduced fractions, yielded rich and dense LC-MS spectra not only for the Fab fraction, but strikingly now also for the Fab2 fraction. Intriguingly, the resulting Lc and Fd repertoires of the reduced Fab2 and Fab fractions looked very alike (Figure S1D and S1E).

This data provided an important clue on why analyzing the intact IgG4 Fab2 samples by LC-MS did not yield interpretable data; likely all IgG4 clones present in serum had undergone comprehensive FAE, and that level of complexity could not be resolved. We explained this enormous complexity through the following concept. If two B cells each produce and secrete an AA and BB IgG4 antibody, Fab-arm exchange will result in the presence of the AA, AB, and BB antibodies in serum. But when hundreds (or even thousands) of different B cells in a human body produce many unique IgG4 antibodies, the number of distinct IgG4 molecules in serum will inflate. In a stochastic model, where each IgG4 half-molecule (HL) could recombine with any other HL, the Fab2 repertoire is massively expanded compared to the corresponding Fab repertoire (k-combinations with repetitions, or 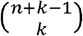). To illustrate this using our data, when analyzing the intact IgG4 Fab profiles (Figure S4A and S4B) we were able to dissect and assign up to 300 clones per serum sample, each having a unique mass and retention time. Assuming a complete and stochastic Fab-arm exchange process, these 300 Fab clones would result in 45,150 distinct bispecific Fab2 molecules. The 300 Fab clones were all high abundant and thus assignable in our LC-MS chromatograms. But when considering the stochastic Fab-arm exchange process, the resulting abundance of each Fab2 molecule would be ∼1000 times diluted. Consequently, intact Fab2 molecules are — because of their enormous diversity and consequently their inherent low abundance — inseparable and undetectable by our current LC-MS based approach.

Upon reduction, however, the complexity originally present in the fully Fab-arm exchanged Fab2 fraction could be brought back to just around 300 Lc and 300 Fd detectable fragments. Similarly, reduction of the Fab fraction would also result in just 300 Lc and 300 Fd fragments. Indeed, by reducing the IgG4 Fab2 and Fab fractions we were able to acquire signal for both the Fab2 and the Fab fractions for all donors, allowing us to compare the IgG4 profiles and clonal repertoires between donors, within a donor over time, and between the clones present in the Fab2 or Fab fractions.

### Fab-arm exchange is a stochastic phenomenon in serum IgG4

Next, we set out to evaluate the differences in IgG4 repertoires obtained from the Fab2 and Fab fraction within the IgG4 serum sample obtained from each individual donor. For this, we had samples from 7 healthy donors available (donor 2 sampled at 7 timepoints). We next analyzed 26 (13/13) paired samples of reduced Fab2 and Fab fractions. Strikingly, within a donor, the repertoires of reduced Fab2 and Fab fractions looked almost identical, both in composition and in fragment abundance (Figure 3A, 3B, Figure S2A, S2B, and S3). This data provides explicit evidence for Fab-arm exchange being completely stochastic *in vivo* with no bias in some IgG4 clones becoming more/less exchanged than others. Additionally, we observed a high degree of overlap between the IgG4 profiles originating from the longitudinal serum samples taken from donor 2, even though these were sampled over a period of more than a year, indicating a long-lasting persistence of the here detected serum IgG4 repertoire (Figure 3C, Figure S3). Of note, the here covered time span is substantially longer than the half-life of IgG4 in circulation (∼21 days^29^). Looking at the data for distinct donors, we observed barely any overlap in detected clones, indicating that IgG4 repertoires are highly personalized and unique.

**Figure 3.**
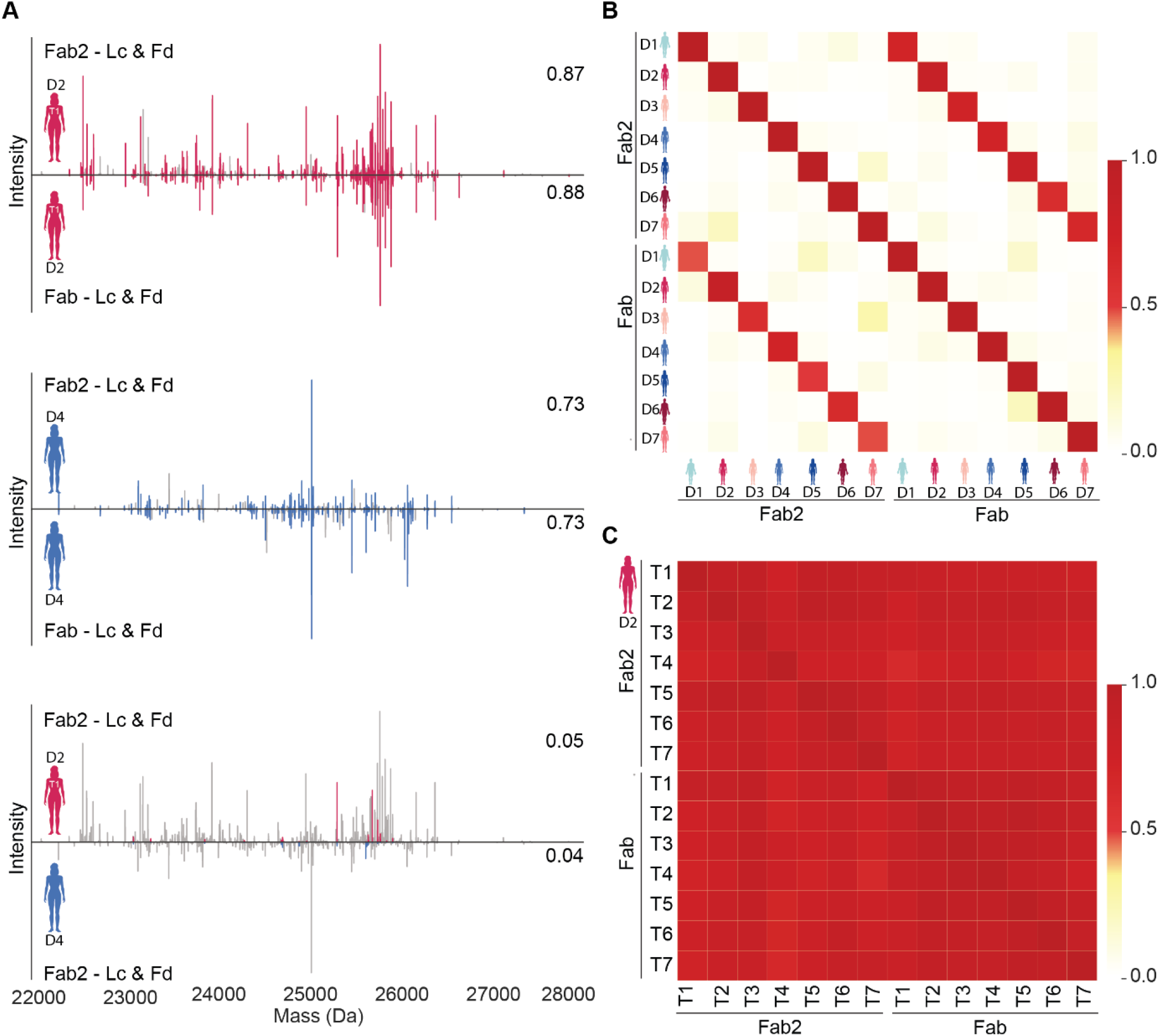
IgG4 repertoire analysis provides unique insights into serum IgG4 compositional properties. A) IgG4 repertoires, displayed as mass plots of the corresponding Fd and Lc fragments (22-28 kDa) generated by reducing the Fab2 (top) or Fab (bottom) fraction. Each line represents a unique Lc or Fd fragment as determined by matching mass and retention time. The fragment abundance is indicated by the height. Compared here are the Fab2 and Fab repertoires from donor 2 (top), donor 4 (middle) and the Fab2 repertoires of donor 2 and 4 (bottom). Repertoires from the same donor are highly similar, whereas repertoires from distinct donors reveal barely any clonal overlap. Shared clonal fragments are colored and unique fragments are depicted in grey. The similarity — based on fragment-intensity overlap between the different mass plots — is indicated on the right corner of each plot. B) Similarity between reduced Fab2 and Fab repertoires from seven distinct individuals. Fab2 and Fab repertoires within a donor are characterized by a high degree of similarity (dark red fields), whereas such a correlation is low when comparing profiles between donors. C) Fab2 and Fab repertoires within a donor (intra-donor) over 16 months (7 sampling points) are characterized by a very high degree of similarity (dark red fields).

Additionally, we extended our methods to profile IgG4 repertoires (Figure S4) at the intact Fab level, arguing that Fab molecules are a good proxy for the total repertoire, while retaining the chain pairing. Our results from the intact Fab profiling are in line with the repertoire-wide observations obtained from the Lc-Fd profiling, validating that IgG4 repertoires are at the Fab-level unique for each individual and dominated by a few hundred high abundant clones. However, at the intact IgG4 level these hundreds of clones will lead to the co-occurrence of tens of thousands of bispecific IgG4 molecules.

## Discussion

In humans four subclasses of immunoglobulin IgG co-exist. Of these four, IgG1 and IgG2 are normally by far the most abundant in serum (with IgG4 being substantially lower)^30^. However, next to IgG1, IgG4 is frequently considered as a platform for applications in biotherapy^31^. Despite its extensive use as biotherapeutics, IgG4 remains a rather understudied IgG subclass, especially at the level of endogenous antibodies^32,33^. Although all four IgG subclasses exhibit structural and functional similarities, they also harbor class-specific features. Here, we address the IgG4-specific feature known as Fab-arm exchange. Because of the more reduction-sensitive IgG4 hinge region and relatively weaker CH3:CH3 interaction dissociation of two half-molecules can occur. Subsequently, half-molecules can either reassemble with their original counterpart, or with a half-molecule from another IgG4 clone. There have been several important pioneering studies investigating this mechanism^8-16^. It is by now well-established that IgG4 Fab-arm exchange is lambda/kappa light-chain independent^16^. Here, we not only extended upon such studies, but also demonstrate for the first time that Fab-arm exchange is independent of the entire Fab-arm. Such insights require simultaneous monitoring of a large endogenous IgG4 population and cannot easily be achieved by studies just focusing on monoclonal IgG4. Furthermore, in IgG4 therapeutics, the underlying natural scaffold is often modified by mutating the serine 228 to a proline, as found in IgG1, to prevent the Fab-arm exchange reaction. While this mutation is useful for regulatory purposes^34^, such antibodies do no longer represent natural IgG4 molecules, and cannot be used as a model in IgG4 research.

Here, we monitor simultaneously the mass and abundance of hundreds of serum IgG4 clones, allowing us for the first time to qualitatively and quantitatively assess the nature and comprehensiveness of Fab-arm exchange. Our initial SEC experiments confirmed already that IgG4 behaves quite distinctively from other IgGs. The digestion of IgGs below the hinge using IdeS produced for all IgGs solely Fab2 fragments, but for IgG4 also a sizeable amount of Fab molecules was observed. This behavior can be explained by the co-occurrence of covalently and non-covalently bound half-molecules of IgG4 (heavy chain and light chain), and is well in line with earlier findings of coexisting full IgG4 and IgG4 half-molecules in recombinant monoclonal IgG4^35-37^.

In the endogenous population of IgG4 in serum, we find that ∼80% of the molecules represent covalently bound full IgG4, making the remainder ∼20% being present as non-covalently bound IgG4. The here reported ratio is at the lower end of previously reported data (> 99% to 75 %)^8,36-38^, but we are confident that our methodology allows for a good estimation of a biologically relevant average as we account for the entire IgG4 repertoire from several individuals rather than single mAb constructs. Additionally, it is interesting to observe how stable the ratios between covalent and non-covalent IgG4 are within and between individuals. We have furthermore validated the capability of our platform to pick up isomeric differences between three monoclonal IgG4, one of which (Natalizumab) has been reported to contain just a limited small fraction of non-covalent isomers^8^.

Based on the apparent absence of detectable high abundant clones when performing LC-MS on the Fab2 fractions, which unmistakably contained protein, and the fact that we could resolve this issue by reducing the Fab2 samples to Lc and Fd fragments, we conclude that Fab-arm exchange affects effectively all IgG4 molecules in serum *in vivo*. As nearly all previous estimates were based on exchange processes involving a recombinant monoclonal IgG4^13,39^, this is the first time that this conclusion is supported by repertoire-wide evidence. Consequently, the repertoire of IgG4 distinctive entities present in blood becomes massively inflated (compared to e.g. IgG1), potentially surpassing even upper estimates for the size of the entire immunoglobulin repertoire of up to one million total circulating immunoglobulins (Figure 4)^40^.

**Figure 4.**
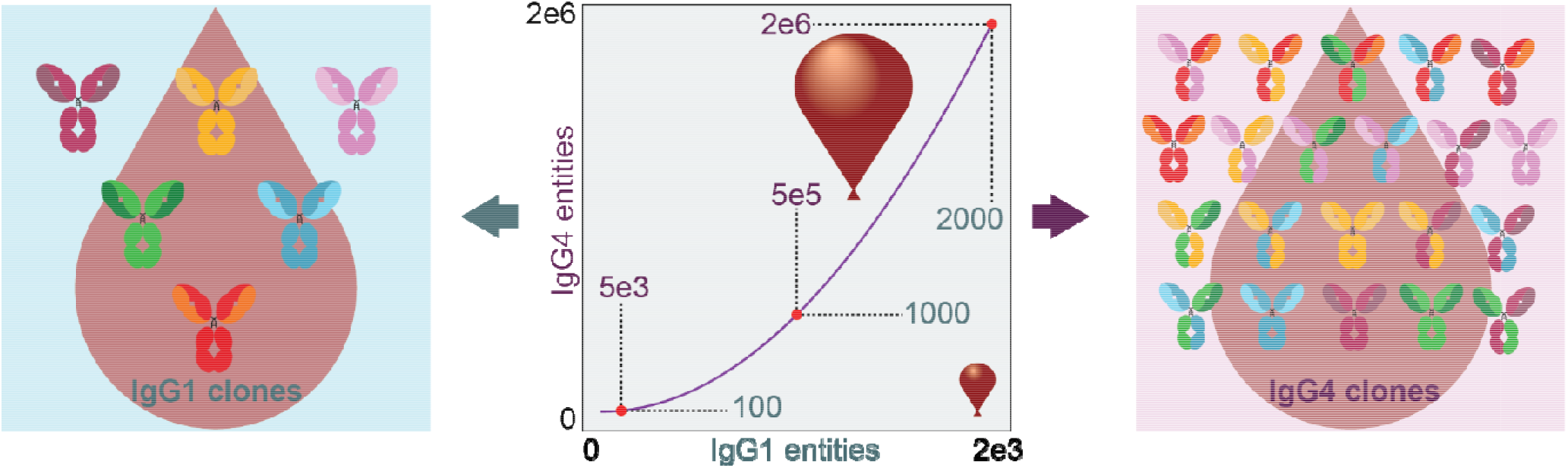
Fab-arm exchange inflates the serum repertoire of IgG4 entities. While all other IgG subclasses (here represented by IgG1, left panel) are secreted as homo-dimers of identical Hc-Lc pairing, IgG4 will experience a complete Fab arm exchange (right panel), producing all possible hetero-dimeric entities as well. If n molecular IgG entities exchange stochastically their Fab arms, it will result in possible combinations. The middle panel displays the massive diversity increase in IgG4 entities (purple values) compared to 100, 1000 and 2000 IgG1 molecular entities (green values). To illustrate further, using a lower estimate of thousand distinctively detected clones in serum for IgG1, this would correspond to more than half a million of distinctive IgG4 entities being present in the same sample.

Within the IgG4 community it has been proposed that the FAE is a stochastic process^33^, although this hypothesis is mostly supported by studies with patient-derived antigen-specific antibodies as well as with monoclonal antibodies^12^. Actual evidence with clonal resolution, however, has not been shown. Here, we further substantiate this prevailing assumption in the field, which is particularly pertinent given that previously analyzed IgG4 monoclonals vary in the proportions of noncovalent hinge isomers, and Fab domains likely influence the course of the hinge isomerization^8^. Yet, if an IgG4 with a lower tendency to form intrachain disulfide bridges takes longer to provide half-molecules for the reaction, this should ultimately not affect the equilibrium of global exchange reactions. Our observations do not exclude the possibility that ‘stability sinks’ of preferred IgG4 half-molecule combinations may exist, but their effect is by no means a dominating factor in IgG4 repertoires of healthy donors as studied here.

In addition to the findings on the structure of endogenous IgG4, we were able to describe expressed IgG4 repertoires with clonal resolution for the first time. Surprisingly, they are dominated by only a few IgG4 clones, which are also unique for an individual and thus represent immunoglobulin fingerprints. Interestingly, our longitudinal observation of a donor suggests that IgG4 repertoires remain stable over time in the absence of an immune challenge. Despite the genomically far distance between IgG4 and IgG1 or IgA1^41^, these observations closely resemble what we have observed before for these immunoglobulin relatives^23,24,42^.

Besides these new insights into IgG4 biology, we have here described a novel approach for studying endogenous IgG4 repertoires. This method could be applied to gain knowledge on the nature of the reported increased IgG4 responses upon mRNA vaccination^43-45^. Furthermore, other research domains may benefit from IgG4 repertoire analysis, such as IgG4 autoimmune diseases (IgG4-AID)^46^ and IgG4-related diseases (IgG4-RD)^47,48^.

Its exceptional ability to exchange half-arms with any other IgG4 makes it indispensable to consider both, structure and composition, when analyzing IgG4 repertoires. Here, we were able to do this, and revealed that this Janus-headed IgG harbors probably the most diverse repertoire of all antibody classes.

## Author contributions

LW, AB and AJRH conceived the project. MGH advised on IgG4 biology and purification of IgG4 from serum. AB and AJRH supervised the study. LW performed all experiments and analyzed the data. All drafted and edited the manuscript. AJRH provided funding for the project.

## Acknowledgements

This research received funding by the Netherlands Organization for Scientific Research (NWO) through CHEMIE.PGT.2023.009 and the Spinoza Award SPI.2017.028 to AJRH. Additionally, continuous support from Genmab (Utrecht, NL) is appreciated, through the gift of recombinant mAbs (team lead Aran Labrijn) and financial support for our work. We thank Arjan Barendregt and Nadia Mokiem for their technical support during the method development and Gestur Vidarsson (Sanquin Research) for providing us with serum samples. We would further like to express our gratitude to Aran Labrijn for critically evaluating the manuscript, and to Oscar Dekker and Jessica van Bokkum (Leiden University Medical Center) for valuable discussions on how best to purify IgG4. MGH receives financial support from the LUMC Gisela Thier 2021 fellowship, PPS Health Holland match call 2021, a Dutch incentive grant and the European Union through a ERC starter grant IgG4-START nr: 101163002.

## Materials and Methods

### Origin of the serum samples and recombinant mAbs

Individual serum samples from healthy donors were provided by Prof. G. Vidarsson from Sanquin Research (Amsterdam, The Netherlands) and obtained in accordance with the ethics board of Sanquin and after informed consent from all donors. The samples were stored at −80 °C until used for processing. For method standardization and development, we used monoclonal antibodies (mAbs) that were provided to us as gifts by Aran Labrijn of Genmab, Utrecht, NL (7D8-IgG1, -IgG2, -IgG3, and -IgG4, and DNP-G2a2-IgG4) or were purchased (Natalizumab) from Evidentic (Potsdam, Germany).

### Serum IgG4 purification and digest

Immunoglobulin enrichment strategies were adapted from or inspired by previously published protocols^24,49^. First, IgG4 were affinity purified from serum using CaptureSelect IgG4 (Hu) affinity matrix (Thermo Fisher Scientific, Leiden, Netherlands). 20 µL bead slurry was added to Pierce spin columns with screw cap (Thermo Fisher Scientific, Rockford, Illinois, USA). The beads were washed three times with 150 µL anticoagulant (375 mM saccharose, 82.5 mM Sodium citrate, 17.5 mM citric acid monohydrate) by centrifugation at 500 x g and room temperature (RT). A plug was inserted to the bottom of the spin column, and 100 µL human serum mixed with 50 µL anticoagulant were added. As an internal standard 200 ng Natalizumab IgG4 was spiked into each sample. The samples were incubated for 1h, 750 rpm at RT on an Eppendorf thermal shaker (Wesseling-Berzdorf, Germany) with a ThermoTop. Afterwards, the plugs were removed from the spin column, and the serum/anticoagulant mixture was collected into an Eppendorf low-bind tube. The IgG4 beads were washed three times with 200 µL PBS (+ 1 M NaCl), two times with 200 µL MilliQ water and subsequently centrifuged for 1 min at 500 x g. IgG4 Fab2 were prepared by in-solution digestion. The Ig were eluted from the matrix into 12 µL phosphate neutralization buffer (429.3 mM sodium dihydrogen phosphate monohydrate, 470.7 disodium hydrogen phosphate dihydrate, pH 7.54) in low-bind Eppendorf tubes by adding 20 µL 0.1 M glycine (pH 2.7) onto the CaptureSelect affinity matrices. After incubation for 5 min, 750 rpm at RT on an Eppendorf thermal shaker, the spin columns were centrifuged at 1000 x g. Two additional steps of adding 20 µL 0.1 M glycine, incubation and centrifugation were performed. Lyophilized FabRICATOR (IdeS, Genovis, Kävlinge, Sweden) was reconstituted in phosphate buffer (PB) (71.55 mM sodium dihydrogen phosphate monohydrate, 78.45 mM disodium hydrogen phosphate dihydrate, pH 7), and 50 U in 50 µL were added to the sample. The sample was digested for 18 h, 750 rpm at 37 °C on an Eppendorf thermal shaker with a ThermoTop. Fc and undigested antibodies were depleted from the sample by performing the Ig specific enrichment as described above. After binding, the spin columns were centrifuged for 1 min at 1,000 x g and IgG4 Fab2 were collected in low-bind Eppendorf tubes.

### Serum IgG1 purification and digest

IgG1 were affinity purified from serum using CaptureSelect IgG1 (Hu) affinity matrix (Thermo Fisher Scientific, Leiden, Netherlands). 20 µL bead slurry was added to Pierce spin columns with screw cap (Thermo Fisher Scientific). The beads were washed three times with 150 µL PBS and centrifuged at 500 x g, RT. A plug was inserted to the bottom of the spin column, and 10 µL human serum mixed with 140 µL PBS were added. As an internal standard 200 ng of IgG1-7D8, IgG2-7D8, IgG3-7D8, and IgG4-7D8 were spiked into the sample to validate specificity. The sample was incubated for 1h, 750 rpm at RT on a Eppendorf thermal shaker with a ThermoTop. The IgG1 beads were washed five times with 200 µL PBS and subsequently centrifuged for 1 min at 500 x g. Lyophilized FabRICATOR was reconstituted in PBS, and 50 U in 50 uL were added to the sample. The sample was digested for 18 h, 750 rpm at 37 °C on an Eppendorf thermal shaker with a ThermoTop. For collection, the spin columns were centrifuged for 1 min at 1,000 x g and IgG1 Fab2 were collected in low-bind Eppendorf tubes.

### Size exclusion chromatography (SEC)

For size exclusion chromatography, an Agilent 1260 Infinity HPLC system (Agilent, Santa Clara, California, USA) was operated in isocratic flow with an attached SEC column (ACQUITY UPLC BEH SEC, Waters, Framingham, Massachusetts, USA, 200 Å, 1.7 µm particle size, 4.6 mm X 300 mm). For separation, a 20-minute separation method was applied (100 mM ammonium acetate, pH 6.9) at a flowrate of 200 µL*min^−1^. Fab2 were collected between 11.6 and 13.8 minutes, and Fab between 13.8 and 16 minutes. Samples were afterwards frozen in liquid nitrogen and lyophilized (Labconco Freezone 2.5 L −80C, Kansas City, Missouri, USA). Each sample was dissolved in a PBS (137 mM sodium chloride, 2.7 mM potassium chloride, 4.77 mM sodium dihydrogen phosphate monohydrate, 5.23 mM disodium hydrogen phosphate dihydrate) volume adjusted to the integrated SEC-UV trace of the respective fraction.

### LC-MS data acquisition

Intact Fab, Fab2, and fragment chain (Lc and Fd) samples were injected into an Orbitrap Exploris 480 (Thermo Fisher Scientific, Bremen, Germany) using a Vanquish Neo UHPLC (Thermo Fisher Scientific, San José, California, USA). For each run 500 ng of total protein was loaded on a MabPac RP analytical column (150 µm x 150 mm, 4 µm particle size). The column compartment was heated to 80 °C. For separation, a 54-minute gradient from 28% to 38% solvent B (0.1% FA, 0.02% DFA in ACN) at a flow rate of 150 μL*min-^1^ was applied to the mobile phase (0.1% FA, 0.02% DFA in ddH_2_O). MS data were collected in Intact Protein and Low Pressure mode. The mass spectrometer was operated in positive ionization mode. The spray voltage was set at 2 kV, the capillary temperature to 275 °C, and the probe heater temperature to 100 °C. The RF Lens was set to 100% (Fab2), 60% (Fab), or 40% (Fd/Lc). Source fragmentation was enabled and set to 15 V (Fab and Lc/Fd), or 50 V (Fab2). Fab and chains samples were recorded using MS1 methods (800 – 3200 m/z for Fab, 600 – 2,400 m/z for Fd and Lc fragment chains) with a maximum injection time (maxIT) of 50 ms, a resolution of 7,500, and a normalized automatic gain control (AGC) target value of 300%. More efficient Fab2 transmission was facilitated by measuring these samples in a targeted MS2 method (scan range 1,000 – 4,000 m/z) with a 1,600 m/z isolation window around 2,400 m/z. HCD collision energy was set to 5 V, AGC target was set to standard, and maximum injection time to Auto settings.

### Clonal profiling data processing and analysis

Fab2, Fab, Lc or Fd masses were obtained by deconvolution from RAW files with BioPharmaFinder 3.2 (Thermo Scientific, San José, California, USA) using the ReSpect algorithm. Spectra between 6 and 60 min were considered and source spectra were generated by the sliding windows algorithm with 0.1 min target average spectrum width, 25% offset, merge tolerance of 30 ppm, and noise rejection set at 95%. The output mass range was set at 20,000 to 65,000 Da for Fab samples, 20,000 to 60,000 Da for reduced samples, and 40,000 to 110,000 Da for Fab2 samples. The target mass was set to 48,000 Da for Fab fractions, 98,000 Da for Fab2 fractions, and 25,000 Da for reduced Lc and Fd fragment samples. Charge states between 10 and 60 (Fab, Lc and Fd), and 15 and 100 (Fab2) were included, respectively, and the Intact protein peak model was selected. RESULTSBPF files were further analyzed using Python 3.10.9 (libraries: Pandas 1.5.3, Numpy 1.23.5, Scipy 1.12.0, Matplotlib 3.7.0, and Seaborn 0.12.2). Component masses generated by BioPharmaFinder were recalculated using an intensity weighted mean considering only the most intense peaks comprising 90% of the total intensity. Components between 46 and 55 kDa (Fab), 92 and 110 kDa (Fab2), and 20 and 30 kDa (Lc and Fd) with the most intense charge state above 1,000 Thomson (Fab2 and Fab) and 700 Thomson (Lc and Fd), respectively, and a score above 40 were considered as derivatives of IgG clones. Clones were matched between runs using average linkage (unweighted pair group method with arithmetic mean UPGMA) L∞ distance hierarchical clustering. Flat clusters were formed based on a cophenetic distance constraint derived from the mass and retention time tolerance. These tolerances were defined as 1.5 Da and 5 minutes for chains and 1.5 Da and 1 minute for Fab. Clones within a flat cluster were considered identical between runs. For intact Fab retention times were aligned based on the spiked in mAb to minimize deviations between runs.

### Clonal filtering

To avoid introducing shared components (e.g., Fc or co-purified serum proteins) all data was rigorously filtered. Components identified in more than two of the seven different donors were removed. We then determined the overlap between reduced and intact samples by matching components based on the above described hierarchical clustering with 5 min and 5 Da tolerance settings. We removed every component that was identified in at least one reduced and unreduced sample, as those were with high likelihood other co-enriched serum proteins. Thereafter, the cleaned two datasets on intact Fab and reduced samples were processed separately. Each of the two datasets underwent a match between run step with 1.5 Da and 1 min tolerance (intact), respectively 1.5 and 5 min tolerance (reduced). We further removed all components with atypical elution behavior > 2 minutes. Contaminants introduced by the breakthrough peak were removed by analyzing only components which were most abundant after 9 (Fab2), respectively 11 minutes (Fab). Lastly, we removed all contaminants identified in three or more of the seven different donors.

