## supplementary information for "Fab-arm exchange affects each and all endogenous serum IgG4 evenly"

**Contains Supplemental Figures 1, 2, 3 and 4, and Supplemental Table 1 and 2.**

### Supplementary Figures

#### *Supplemental Figure 1*

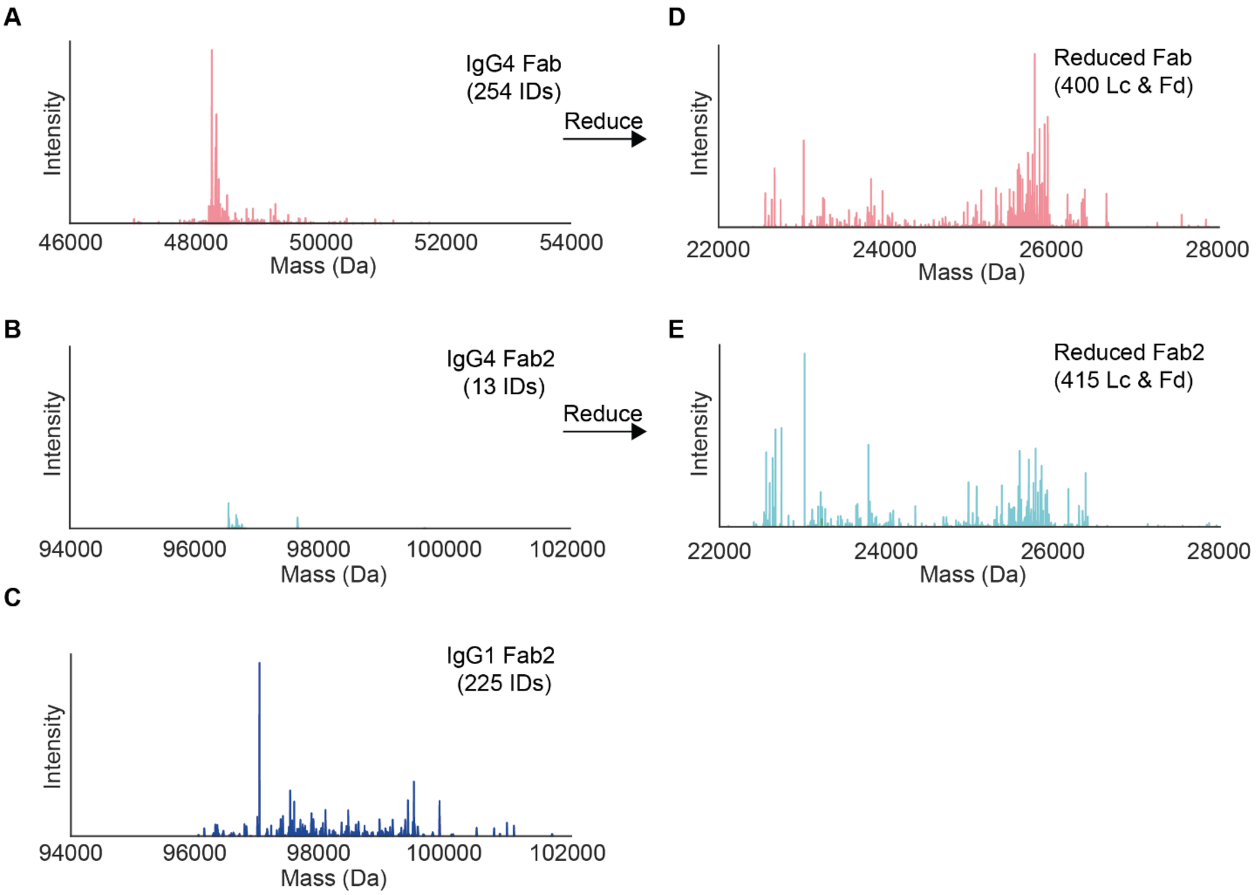


**Figure S1 | Profiling of IgG4 repertoires requires dedicated workflows**. *Provided are mass plots of by intact mass LC-MS detected IgG4 fragments of IgG4, namely Fab (A), Fab2 (B) and reduced Lc and Fd fragments (D &E), with in (C) Fab2 fragments of IgG1. Each line in the mass plots represents a distinct antibody fragment (Fab, Fab2 or thereof derived reduced fragment chains: Lc and Fd). The abundance is indicated by the height, and the total number of respective studied fragment molecules (IDs) is indicated in the top right corner of each plot. All analyses were performed on the same serum sample (D2T2). A, B, D and E depict measurements of the same serum IgG4 sample under non-reducing (A, B) or reducing (D, E) conditions. The presented mass range is adjusted for the respective IgG4 fragment masses. Illustratively, A) displays a dense clonal IgG4 Fab repertoire (254 clones). B) The corresponding clonal IgG4 Fab2 repertoire is sparsely populated with just over a dozen of detectable Fab2 clones. In contrast in C) is shown that the clonal IgG1 Fab2 repertoire is readily detectable and densely populated (225 clones). In D) is the reduced counterpart of A) depicted and displays a total of 400 fragments (either Lc or Fd). E) represents the reduced counterpart of B) and exhibits contrary to B) a densely populated fragment repertoire (415 Lc or Fd fragments), quite alike to D).*

#### ***Supplemental Figure 2***

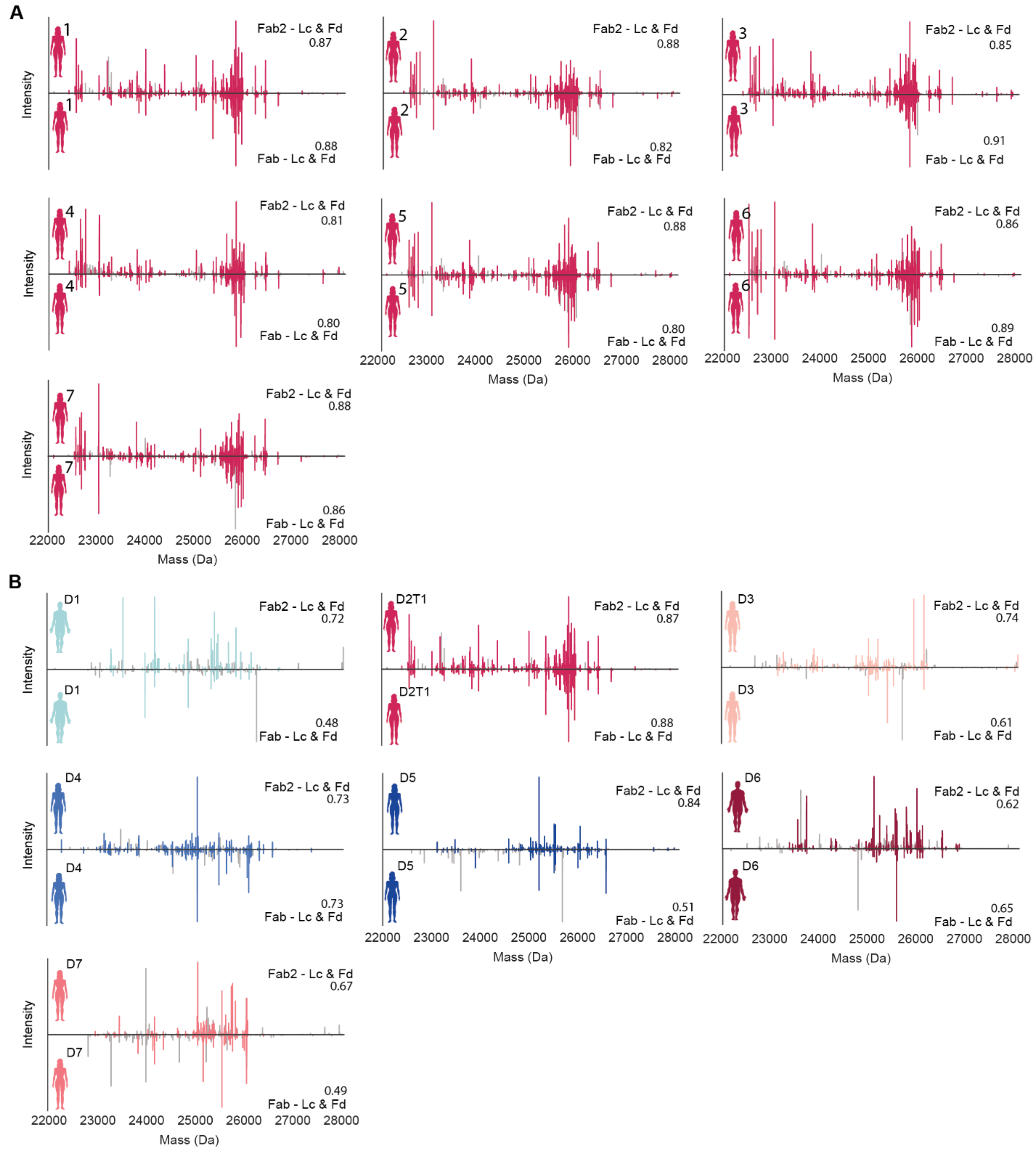


**Figure S2 | Mirror-plots of reduced Lc and/or Fd (22-28 kDa) repertoires originating from either Fab2 (top) or Fab fractions (bottom).** *The observed repertoire similarity is illustrated by the mirrored mass plots in which each line represents a unique fragment at its respective mass and the height indicates the abundance as determined by LC-MS. Unique fragments are characterized by matching mass and retention times. Each mirror plot compares the similarity between the Fab2s and Fabs fractions originating from the same IgG4 serum sample. Cartoons on the left side indicate from which donor a given repertoire originates. Fab2 fractions are displayed at the top of each plot, Fab fractions at the bottom. The numbers on the right side of each plot quantify the similarity, based on intensity-based overlap. A) displays the comparisons of seven (T1-T7) longitudinal serum samples of donor 2, collected over a 16 months' time window . B) displays the comparisons of paired samples from seven healthy donors. All comparisons between Fab2 and corresponding Fab fraction exhibit a high degree of similarity (see also Figure S3).*

#### *Supplemental Figure 3*

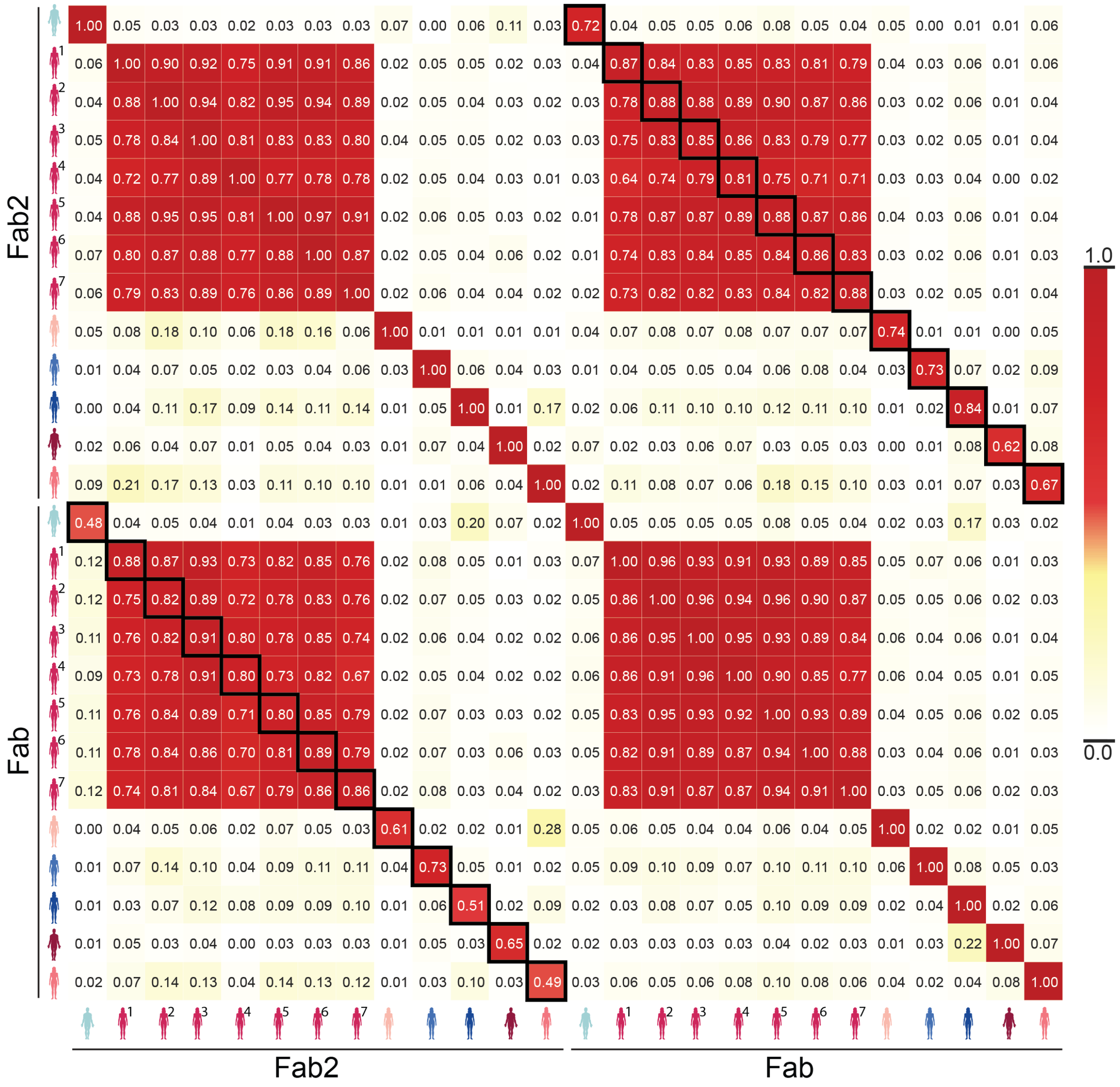


**Figure S3 | Inter- and intra-donor similarities in IgG4 repertoires, comparing the repertoires obtained by analyzing the Lc and Fd fragments originating from the Fab2 and Fab fractions from seven healthy donors.** *For donor 2 seven (T1-T7) longitudinal serum samples were analyzed, collected over a 16 months' time window allowing to assess intra-donor variability. The similarity, based on the intensity-based overlap between the repertoires is indicated by the number in each field using the heat-map ruler on the right. Black squares highlight the pairwise comparisons between corresponding Fab2 and Fab fractions of exactly the same serum sample. Repertoires derived from different donors display low (inter-donor) similarity, both for the reduced Fab and Fab2 fraction (light colored fields). Contrary, repertoires from the same donor display a high degree of overlap over time and between the reduced fractions (black squares, dark red fields). High inter-donor similarity is also seen when comparing Fab and Fab2 repertoires of the longitudinal samples within and between each other (large clusters of red fields), hinting at a high degree of IgG4 clonal repertoire stability over time in this donor 2.*

#### Supplemental Figure 4

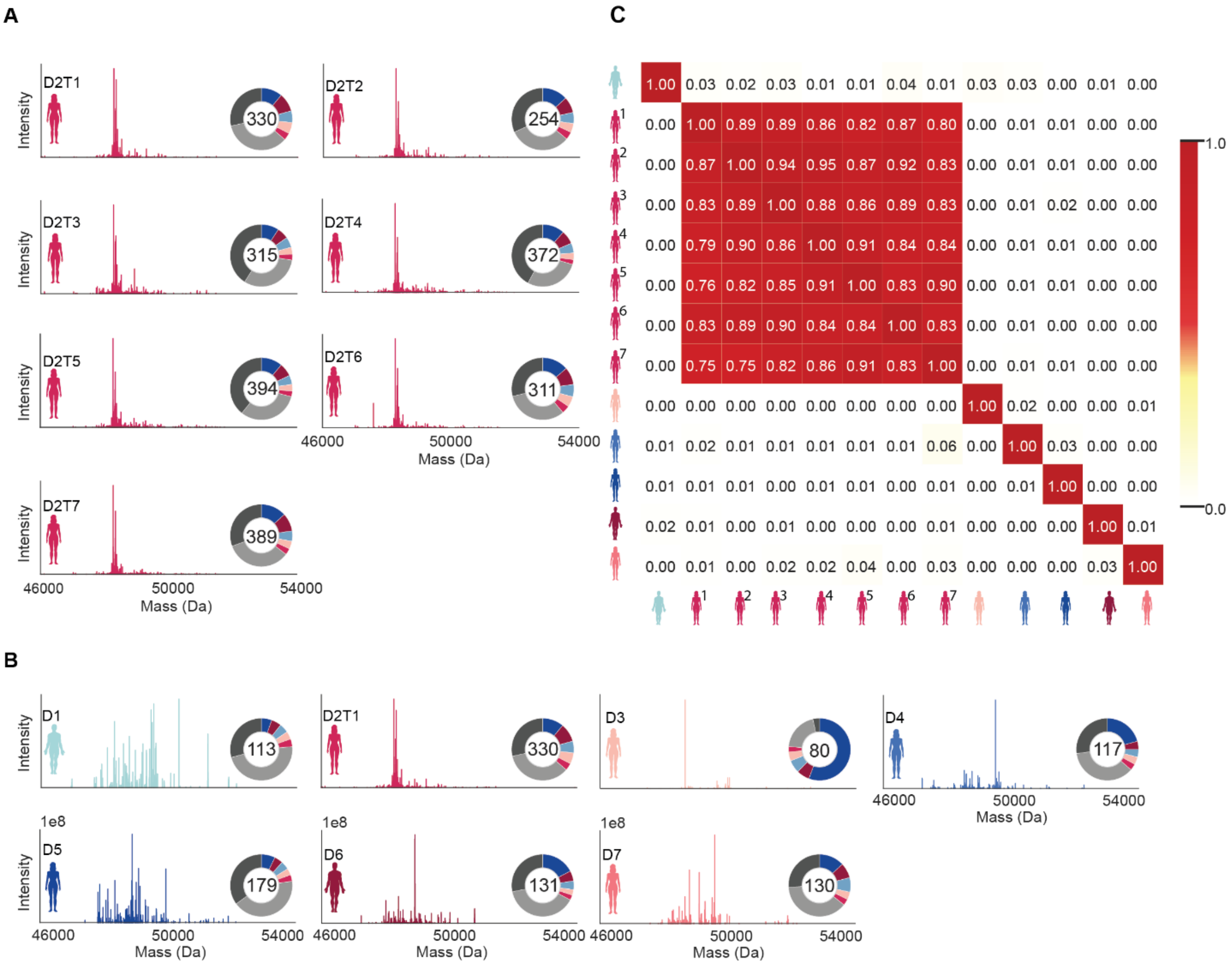


**Figure S4 | Inter- and intra-donor similarities in IgG4 repertoires, comparing directly the Fab repertoires from the seven healthy donors.** *Clonal abundance is indicated by line height. The donor corresponding to the repertoires is named in the top-left corner of each mass plot. The pie charts on the right side of each plot display the total number of identified clones in the middle (range 80 – 394), and the detected clonal dispersity. Color-depicted are the contributions of the top 5 highest abundant Fab clones to the total detected Fab intensity, in light-grey the contributions of the top 6-30, and in dark-grey the contribution of the residual clones. A) displays the longitudinal clonal IgG4 Fab repertoires of healthy donor 2. B) displays the clonal IgG4 Fab repertoires of seven healthy donors. The mass plots reveal that IgG4 repertoires from the same donor look very similar, whereas repertoires from different donors are highly divergent. The top 30 highest clones are responsible for about two third of the total clonal intensity detected in each experiment. C) Heatmap depicting the pairwise comparison of clonal IgG4 repertoires. The corresponding donors are indicated by the cartoons on the axes using the same color scheme as in A) and B). The numbers in the fields show the similarity based on clone-intensity overlap between the Fab-based repertoires. A high degree of overlap is indicated by dark red fields, low overlap by light-colored fields. While repertoires from different donors show barely any similarity, repertoires derived from an individual donor retain very much alike.*

### Supplementary Tables

#### Supplemental Table S1

**Supplementary Table S1: Fab2 and Fab proportions in each analyzed serum and recombinant IgG4 sample as determined by UV-SEC.** *Shown are the values obtained for serum IgG4 (upper two panels) and three monoclonal IgG4's with corresponding means and standard deviations (lower two panels). The proportions were calculated by integrating the area under the curve between 11.6 and 13.8 minutes (Fab), respectively 13.8 and 16 minutes (Fab2). One time point of donor 2 (D2T1) was excluded as the IgG4 yield was only sufficient for preparative, but not analytical SEC. The donors used to calculate the serum IgG4 mean are indicated in black, additional timepoints of donor 2 in grey.*

| Sample | Proportion Fab | Proportion Fab2 |
|---|---|---|
| | ***Serum IgG4*** | |
| D1 | 29.3 | 70.7 |
| D2T2 | 16 | 84 |
| D2T3 | 16.1 | 83.9 |
| D2T4 | 17.1 | 82.9 |
| D2T5 | 17 | 83 |
| D2T6 | 17.4 | 82.6 |
| D2T7 | 16.5 | 83.5 |
| D3 | 23.7 | 76.3 |
| D4 | 19.6 | 80.4 |
| D5 | 20.7 | 79.3 |
| D6 | 24.5 | 75.5 |
| D7 | 23.2 | 76.8 |
| | ***across all serum IgG4*** | |
| Mean (all samples) | 20.1 | 79.1 |
| Mean seven donors | 22.4 | 77.6 |
| Standard deviation seven donors (SD) | 3.9 | 3.9 |

| Sample | Proportion Fab | Proportion Fab2 |
|---|---|---|
| | ***Monoclonal IgG4*** | |
| 7D8 | 17.1 | 82.9 |
| Natalizumab | 4.7 | 95.3 |
| DNP-G2a2 | 14.6 | 85.4 |
| | ***across all monoclonal IgG4's*** | |
| Mean | 12.1 | 87.9 |
| Standard deviation (SD) | 5.4 | 5.4 |

#### Supplemental Table S2

**Supplemental Table S2:** *Overview of selected properties of used monoclonal antibodies. Amino acid sequences of chains from the used monoclonal antibodies (mAbs), corresponding fragment masses and observed post translational modifications. The light grey portions in the Hc column correspond to the cleaved Fc.*

| mAb | Heavy chain (Hc) | Light chain (Lc) | Fab2 mass<br>Fab mass<br>Fd mass<br>Lc mass | Modification |
|---|---|---|---|---|
| **Natalizumab** | QVQLVQSGAEVKKPGASVKVSCKASGFNIKDTYIHWVRQAPGQR<br>LEWMGRIDPANGYTKYDPKFQGRVTITADTSASTAYMELSSLRSE<br>DTAVYYCAREGYYGNYGVYAMDYWGQGTLVTVSSASTKGPSVFP<br>LAPCSRSTSESTAALGCLVKDYFPEPVTVSWNSGALTSGVHTFPAV<br>LQSSGLYSLSSVVTVPSSSLGTKTYTCNVDHKPSNTKVDKRVESKY<br>GPPCPSCPAPEFLGGPSVFLFPPKPKDTLMISRTPEVTCVVVDVSQ<br>EDPEVQFNWYVDGVEVHNAKTKPREEQFNSTYRVVSVLTVLHQ<br>DWLNGKEYKCKVSNKGLPSSIEKTISKAKGQPREPQVYTLPPSQEE<br>MTKNQVSLTCLVKGFYPSDIAVEWESNGQPENNYKTTPPVLDSD<br>GSFFLYSRLTVDKSRWQEGNVFSCSVMHEALHNHYTQKSLSLSLG<br>K | DIQMTQSPSSLSASVGDRVTITCKTSQDINKYMAWYQQTPGKAPR<br>LLIHYTSALQPGIPSRFSGSGSGRDYTFTISSLQPEDIATYYCLQYDNL<br>WTFGQGTKVEIKRTVAAPSVFIFPPSDEQLKSGTASVVCLLNNFYPR<br>EAKVQWKVDNALQSGNSQESVTEQDSKDSTYSLSSTLTLSKADYEK<br>HKVYACEVTHQGLSSPVTKSFNRGEC | 98,418.0 Da<br>49,209.0 Da<br>25,664.8 Da<br>23,556.3 Da | Pyroglutamate (Hc) |
| **7D8-IgG1** | EVQLVESGGGLVQPDRSLRLSCAASGFTFHDYAMHWVRQAPGK<br>GLEWVSTISWNSGTIGYADSVKGRFTISRDNAKNSLYLQMNSLRA<br>EDTALYYCAKDIQYGNYYYGMDVWGQGTTVTVSSASTKGPSVFPL<br>APSSKSTSGGTAALGCLVKDYFPEPVTVSWNSGALTSGVHTFPAVL<br>QSSGLYSLSSVVTVPSSSLGTQTYICNVNHKPSNTKVDKRVEPKSC<br>DKTHTCPPCPAPELLGGPSVFLFPPKPKDTLMISRTPEVTCVVVDV<br>SHEDPEVKFNWYVDGVEVHNAKTKPREEQYNSTYRVVSVLTVLH<br>QDWLNGKEYKCKVSNKALPAPIEKTISKAKGQPREPQVYTLPPSRE<br>EMTKNQVSLTCLVKGFYPSDIAVEWESNGQPENNYKTTPPVLDSD<br>GSFFLYSKLTVDKSRWQQGNVFSCSVMHEALHNHYTQKSLSLSPG<br>K | EIVLTQSPATLSLSPGERATLSCRASQSVSSYLAWYQQKPGQAPRLLI<br>YDASNRATGIPARFSGSGSGTDFTLTISSLEPEDFAVYYCQQRSNWPI<br>TFGQGTRLEIKRTVAAPSVFIFPPSDEQLKSGTASVVCLLNNFYPREA<br>KVQWKVDNALQSGNSQESVTEQDSKDSTYSLSSTLTLSKADYEKHK<br>VYACEVTHQGLSSPVTKSFNRGEC | 98,429.9 Da<br>49,215.0 Da<br>25,785.0<br>23,442.1 | |
| **7D8-IgG2** | EVQLVESGGGLVQPDRSLRLSCAASGFTFHDYAMHWVRQAPGK<br>GLEWVSTISWNSGTIGYADSVKGRFTISRDNAKNSLYLQMNSLRA<br>EDTALYYCAKDIQYGNYYYGMDVWGQGTTVTVSSASTKGPSVFPL<br>APCSRSTSESTAALGCLVKDYFPEPVTVSWNSGALTSGVHTFPAVL<br>QSSGLYSLSSVVTVPSSNFGTQTYTCNVDHKPSNTKVDKTVERKCC<br>VECPPCPAPPVAGPSVFLFPPKPKDTLMISRTPEVTCVVVDVSHED<br>PEVQFNWYVDGVEVHNAKTKPREEQFNSTFRVVSVLTVVHQDW<br>LNGKEYKCKVSNKGLPAPIEKTISKTKGQPREPQVYTLPPSREEMT<br>KNQVSLTCLVKGFYPSDIAVEWESNGQPENNYKTTPPMLDSDGSF<br>FLYSKLTVDKSRWQQGNVFSCSVMHEALHNHYTQKSLSLSPGK | EIVLTQSPATLSLSPGERATLSCRASQSVSSYLAWYQQKPGQAPRLLI<br>YDASNRATGIPARFSGSGSGTDFTLTISSLEPEDFAVYYCQQRSNWPI<br>TFGQGTRLEIKRTVAAPSVFIFPPSDEQLKSGTASVVCLLNNFYPREA<br>KVQWKVDNALQSGNSQESVTEQDSKDSTYSLSSTLTLSKADYEKHK<br>VYACEVTHQGLSSPVTKSFNRGEC | 97,859.2<br>48,929.6<br>25,501.6<br>23,442.1 | |
| **7D8-IgG3** | EVQLVESGGGLVQPDRSLRLSCAASGFTFHDYAMHWVRQAPGK<br>GLEWVSTISWNSGTIGYADSVKGRFTISRDNAKNSLYLQMNSLRA<br>EDTALYYCAKDIQYGNYYYGMDVWGQGTTVTVSSASTKGPSVFPL<br>APCSRSTSGGTAALGCLVKDYFPEPVTVSWNSGALTSGVHTFPAVL<br>QSSGLYSLSSVVTVPSSSLGTQTYTCNVNHKPSNTKVDKRVELKTP<br>LGDTTHTCPRCPEPKSCDTPPPCPRCPEPKSCDTPPPCPRCPEPKS | EIVLTQSPATLSLSPGERATLSCRASQSVSSYLAWYQQKPGQAPRLLI<br>YDASNRATGIPARFSGSGSGTDFTLTISSLEPEDFAVYYCQQRSNWPI<br>TFGQGTRLEIKRTVAAPSVFIFPPSDEQLKSGTASVVCLLNNFYPREA<br>KVQWKVDNALQSGNSQESVTEQDSKDSTYSLSSTLTLSKADYEKHK<br>VYACEVTHQGLSSPVTKSFNRGEC | 108,581.5<br>54,292.8<br>30,869.8<br>23,442.1 | |

| | | | |
|---|---|---|---|
| | CDTPPPCPRCPAPELLGGPSVFLFPPKPKDTLMISRTPEVTCVVVD<br>VSHEDPEVQFKWYVDGVEVHNAKTKPREEQYNSTFRVVSVLTVL<br>HQDWLNGKEYKCKVSNKALPAPIEKTISKTKGQPREPQVYTLPPSR<br>EEMTKNQVSLTCLVKGFYPSDIAVEWESSGQPENNYNTTPPMLD<br>SDGSFFLYSKLTVDKSRWQQGNIFSCSVMHEALHNRFTQKSLSLSP<br>GK | | |
| **7D8-IgG4** | EVQLVESGGGLVQPDRSLRLSCAASGFTFHDYAMHWVRQAPGK<br>GLEWVSTISWNSGTIGYADSVKGRFTISRDNAKNSLYLQMNSLRA<br>EDTALYYCAKDIQYGNYYYGMDVWGQGTTVTVSSASTKGPSVFPL<br>APCSRSTSESTAALGCLVKDYFPEPVTVSWNSGALTSGVHTFPAVL<br>QSSGLYSLSSVVTVPSSSLGTKTYTCNVDHKPSNTKVDKRVESKYG<br>PPCPSCPAPEFLGGPSVFLFPPKPKDTLMISRTPEVTCVVVDVSQE<br>DPEVQFNWYVDGVEVHNAKTKPREEQFNSTYRVVSVLTVLHQD<br>WLNGKEYKCKVSNKGLPSSIEKTISKAKGQPREPQVYTLPPSQEE<br>MTKNQVSLTCLVKGFYPSDIAVEWESNGQPENNYKTTPPVLDSD<br>GSFFLYSRLTVDKSRWQEGNVFSCSVMHEALHNHYTQKSLSLSLG<br>K | EIVLTQSPATLSLSPGERATLSCRASQSVSSYLAWYQQKPGQAPRLLI<br>YDASNRATGIPARFSGSGSGTDFTLTISSLEPEDFAVYYCQQRSNWPI<br>TFGQGTRLEIKRTVAAPSVFIFPPSDEQLKSGTASVVCLLNNFYPREA<br>KVQWKVDNALQSGNSQESVTEQDSKDSTYSLSSTLTLSKADYEKHK<br>VYACEVTHQGLSSPVTKSFNRGEC | 98,011.2<br>49,005.7<br>25,575.7<br>23,442.1 |
| **DNP-G2a2-IgG4** | DVRLQESGPGLVKPSQSLSLTCSVTGYSITNSYYWNWIRQFPGNKL<br>EWMVYIGYDGSNNYNPSLKNRISITRDTSKNQFFLKLNSVTTEDT<br>ATYYCARATYYGNYRGFAYWGQGTLVTVSAASTKGPSVFPLAPCS<br>RSTSESTAALGCLVKDYFPEPVTVSWNSGALTSGVHTFPAVLQSSG<br>LYSLSSVVTVPSSSLGTKTYTCNVDHKPSNTKVDKRVESKYGPPCPS<br>CPAPEFLGGPSVFLFPPKPKDTLMISRTPEVTCVVVDVSQEDPEVQ<br>FNWYVDGVEVHNAKTKPREEQFNSTYRVVSVLTVLHQDWLNGK<br>EYKCKVSNKGLPSSIEKTISKAKGQPREPQVYTLPPSQEEMTKNQV<br>SLTCLVKGFYPSDIAVEWESNGQPENNYKTTPPVLDSDGSFFLYSRL<br>TVDKSRWQEGNVFSCSVMHEALHNHYTQKSLSLSLGK | DIRMTQTTSSLSASLGDRVTISCRASQDISNYLNWYQQKPDGTVKL<br>LIYYTSRLHSGVPSRFSGSGSGTDYSLTISNLEQEDIATYFCQQGNTLP<br>WTFGGGTKLEIKRTVAAPSVFIFPPSDEQLKSGTASVVCLLNNFYPR<br>EAKVQWKVDNALQSGNSQESVTEQDSKDSTYSLSSTLTLSKADYEK<br>HKVYACEVTHQGLSSPVTKSFNRGEC | 98,782.0<br>49,391.0<br>25,830.0<br>23,573.2 |
